# Development and optimisation of cationic lipid nanoparticles for mRNA delivery

**DOI:** 10.1101/2023.02.07.524134

**Authors:** Dongnan Yan, Haonan Lu, Apanpreet Kaur, Ruisi Fu, Ning Wang, Jin Hui Teh, Hantao Lou, Eric O Aboagye, Rongjun Chen

## Abstract

Messenger RNA (mRNA) has been proposed as a therapeutic agent for various diseases, including cancer. To ensure effective transfection of cancer cells, mRNA needs to be transported with a delivery system that protects its integrity and functionality. In this regard, cationic lipid nanoparticles composed of dioleoylphosphatidylethanolamine (DOPE) and 3β-[N-(N’,N’-dimethylaminoethane)-carbamoyl] cholesterol (DC-Chol) have emerged as common vectors to deliver mRNA. In this project, we aim to use luciferase mRNA as a reporter to synthesise mRNA-loaded cationic lipid nanoparticles, and optimise their mRNA encapsulation and transfection efficiency in ovarian cancer cells. The optimisation process included: 1) adjusting the lipid formulation; 2) adjusting the input mRNA concentration before lipid nanoparticle extrusion; and 3) adjusting the extrusion methods. After optimisation, the encapsulation efficiency was optimised to 62%, thus achieving a relatively high transfection luminescence signal (9.4 times compared to baseline). The lipid nanoparticles also demonstrated stable physical characteristics and high biocompatibility (above 75% cell viability after treatment) within 24 hours. Overall, this project evaluated the synthesis of DOPE/DC-Chol cationic lipid nanoparticles, and optimised their mRNA encapsulation and transfection efficiency in ovarian cancer cell lines. The optimised lipid nanoparticles can be utilised as an ideal system for mRNA delivery, which could be further developed as a potential platform for the immunotherapy in ovarian cancer.

## Background

Messenger RNA (mRNA) is a transient intermediate between DNA and proteins (*1*). *In vitro*-transcribed (IVT) mRNA was created during the late 1980s, owing to research into its structure and functions (*2*). Numerous methods have been investigated to reduce the instability and immunogenicity of IVT mRNA since the original proof-of-concept animal research in 1990 (*3*), among which the improvement of drug delivery technologies has significantly accelerated the preclinical development of mRNA treatments (*4*–*7*).

To fully achieve their therapeutic potency, mRNA molecules need to travel to the target cells and produce enough corresponding proteins. In this case, the delivery systems need to be designed to ensure protection against extracellular RNase breakdown, and concurrently facilitate cellular absorption and endosomal escape of mRNA (*2*, *5*, *8*). In recent years, highly efficient mRNA delivery systems have laid the groundwork for mRNA to act as a new medication (*9*–*11*).

Lipid-based nanoparticles, including lipoplex (*12*), liposomes (*13*), and lipid-polymer hybrid nanoparticles (*14*), have been widely applied to efficient mRNA transport due to their excellent structural flexibility to be labelled with targeting, imaging, and therapeutic compounds (*15*). For example, lipid nanoparticle mRNA vaccines are now being used in clinical trials to treat coronavirus disease 2019 (COVID-19), which is a significant milestone for mRNA treatments (*16*, *17*). Lipid nanoparticles have also been successfully applied for CRISPR/Cas9 delivery (*18*–*20*). Apart from viral vaccines and genome editing, mRNA-loaded lipid nanoparticles have also demonstrated great potential in cancer immunotherapy (*21*, *22*). For example, Oberli et al. have developed an ovalbumin (OVA) mRNA-loaded lipid nanoparticle library to transfect and activate CD8^+^ T cells and dendritic cells, enhancing the immune response in melanoma (*23*). Similar research has also been conducted on pancreatic cancer (*24*, *25*) and lung cancer (*26*).

Several different lipid formulations have been analysed in synthesising lipid nanoparticles (LNPs) (*27*–*29*), among which cationic LNPs were the earliest delivery system that successfully introduced mRNA molecules into target cells (*30*). The electrostatic interaction between positively charged lipids and negatively charged mRNA phosphate backbone can enhance mRNA binding, thus improving the encapsulation efficiency of mRNA (*31*). In addition, cationic lipids can facilitate the interaction with the anionic phospholipids in the plasma membrane and subsequently promote their uptake by endocytosis, demonstrating high *in vitro* transfection efficiency (*32*).

Many cationic and ionisable lipids have been utilised to synthesise mRNA-loaded LNPs, such as 2-di-O-octadecenyl-3-trimethylammonium-propane **(DOTMA)** (*28*), 1,2-dioleoyl-3-trimethylammonium-propane (18:1 TAP, **DOTAP**) (*33*, *34*) and 3β-[N-(N’,N’-dimethylaminoethane)-carbamoyl]cholesterol **(DC-Chol)** (*35*, *36*). In this project, we included **DOTAP** and **DC-Chol** in the nanoparticle formulation because they have higher short-term stability than DOTMA (*37*).

DOTAP is one of the most widely used cationic lipids for gene transfection (*34*). DOTAP has a single quaternary amine **(Figure 1A)** that is positively charged at all physiological pH, so there is one positive charge per molecule. LNPs utilising DOTAP have shown effectiveness for both *in vitro* and *in vivo* transfection (*33*, *34*, *38*). DC-Chol is an ionisable cholesterol derivative (*31*). Cholesterol has been proven to increase LNP stability and retention time (*39*). As cholesterol constitutes around 40% of the mammalian plasma membrane, utilising cholesterol or cholesterol derivatives in lipid formulation could increase the cellular uptake of the LNPs (*40*). DC-Chol·HCl, which contains one protonated nitrogen atom **(Figure 1B)**, has been widely used to produce DC-Chol-related LNPs (*41*). The pKa of the tertiary amine in DC-Cholis around 7.8 (*42*), so it is protonated at physiological pH.

**Figure 1.**
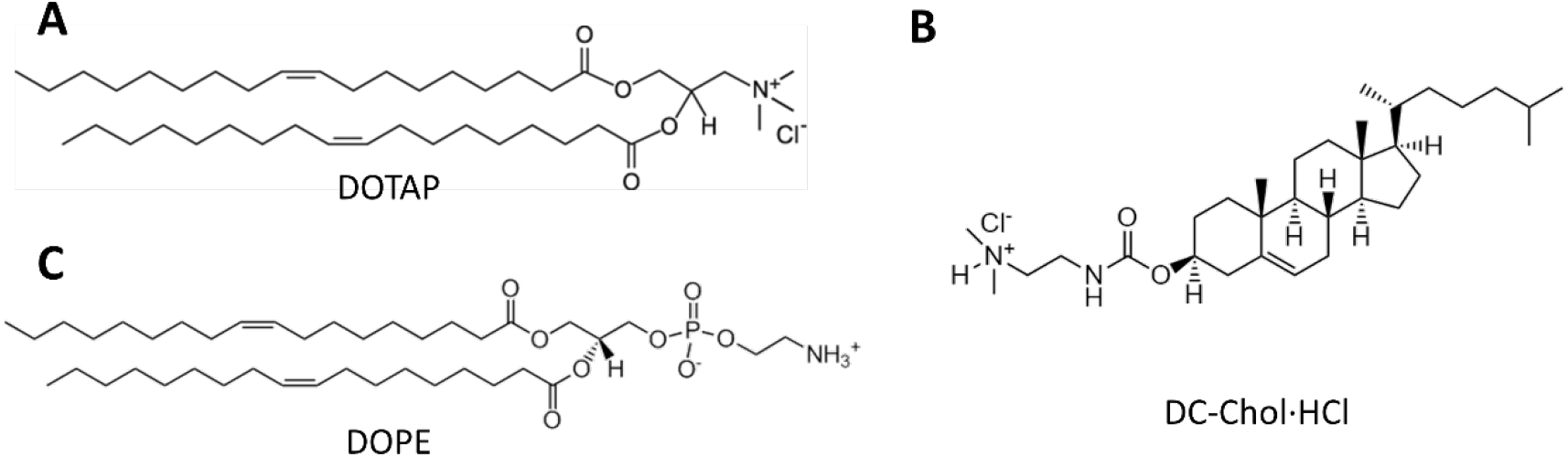
The chemical structures of the lipids and lipid derivatives. **(A)** 1,2-dioleoyl-3-trïmethylammonium-propane (DOTAP). **(B)** 3β-[N-(N’,N’-dimethylaminoethane)-carbamoyl]cholesterol (DC-Chol). **(C)** 1,2-dioleoyl-sn-glycero-3-phosphoethanolamine (18:1 (Δ9-Cis) PE, DOPE).

Despite the high mRNA encapsulation and transfection efficiencies of cationic LNPs, cytotoxicity has become one of the main challenges that limit their applications (*43*). Therefore, involving other helper lipids is essential so as to adjust the biocompatibility of the delivery systems (*44*). For example, 1,2-dioleoyl-sn-glycero-3-phosphoethanolamine (18:1 (Δ9-Cis) PE, DOPE) **(Figure 1C)** is an electricallyneutral helper lipid that could be combined to cationic phospholipids to modify the surface charge of the lipid nanoparticles. It can also increase the effectiveness of RNA transfection by facilitating RNA endosomal escape (*45*).

In recent years, various LNPs have been developed to deliver mRNA to cancer cells, most of which involve DOPE, DOTAP and DC-Chol in the lipid formulation (*28*, *29*, *34*, *36*).

However, the mRNA encapsulation and delivery efficiency of these nanoparticles remain inadequate (*31*). Therefore, further synthetic adjustments are needed to optimise the delivery efficiency of the LNPs.

Various parameters, such as size distribution and zeta potential, can affect the mRNA encapsulation and transfection efficiency of the cationic LNPs (*46*, *47*). Size distribution can affect the transfection efficiency significantly. The diameter of LNPs applied to mRNA transfection is usually limited to less than 150 nm to ensure adequate cellular uptake (*48*). Research has shown that the size of LNPs is negatively correlated with transfection efficiency (*49*). Size distribution can reflect the homogeneity of the LNPs, which can be quantified using Polydispersity Index (PDI) (*50*). Zeta potential represents the overall charge of the particle in a particular medium, which is also an important factor influencing encapsulation and transfection efficiency. Research has shown that zeta potential is positively correlated to the cell-LNP binding, uptake and fusion (*51*).

These parameters can all be optimised by adjusting the ratio of cationic/ionisable lipids in the lipid formulation, input mRNA concentration, and the extrusion process, during which the LNP samples are extruded through a polycarbonate (PC) membrane with a specific pore diameter.

Our lab has recently identified several immune-regulating genes, including IL15 and CXCL10, which are closely related to the immunotherapy response of ovarian cancer. As a starting point, this project aimed to use luciferase mRNA as a reporter gene to synthesise mRNA-loaded LNPs which can efficiently transfect ovarian cancer cell lines. The encapsulation and transfection efficiency of LNPs were optimised by adjusting reaction parameters during the synthesis, including lipid formulation, input mRNA concentration and extrusion process. Apart from optimising encapsulation and transfection efficiency, the stability and biocompatibility of the LNPs were also analysed in a panel of ovarian cancer cell lines.

## Material and Methods

### mRNA Transcription and Phenol Extraction

RiboMAX™ Large Scale RNA Production Systems T7 (Promega, UK) was used to transcribe luciferase mRNA from the linear luciferase DNA template (1,800 bases, Promega, UK). Generally, T7 transcription buffer, T7 enzyme mix, rNTPs (25 mM ATP, CTP, GTP, UTP), and 10 μg of linear DNA template from the kit were mixed and incubated at 37 °C for 4 hours.

After the transcription, RQ1 RNase-free DNase (Promega, UK) was added to the reaction and incubated at 37 °C for 15 minutes to eliminate the DNA template. The mRNA was extracted by citrate-saturated phenol (pH 4.7): chloroform: isoamyl alcohol (125:24:1) (Sigma-Aldrich, UK). After centrifugation at 10,000x g, the upper aqueous phase was isolated, mixed with chloroform: isoamyl alcohol (24:1) (Sigma-Aldrich, UK) and centrifuged again. The upper aqueous phase was then separated, mixed with 0.1 volume of 3M Sodium Acetate (Promega, UK) and 1 volume of isopropanol (Sigma-Aldrich, UK), and placed on ice for 5 minutes. The reaction was centrifuged again, and the resulting RNA precipitation was washed with 70% ethanol and resuspended in RNase-free water.

The concentration of mRNA was then measured by Nanodrop™ 2000 (ThermoFisher, UK).

### Poly(A) polymerase tailing and m^7^G capping of mRNA

The m^7^G capping was carried out according to the protocol of the ScriptCap™ m^7^G Capping System (CellScript, UK). Generally, the purified RNA was denatured at 65 °C for 10 minutes and mixed with 10x ScriptCap capping buffer, 10 mM GTP, 2 mM S-adenosyl-methionine (SAM), ScriptGuard RNase inhibitor and ScriptCap capping enzyme. The mix was then incubated at 37 °C for 30 minutes.

Following the m^7^G capping, the poly(A) polymerase tailing was carried out according to the protocol of the A-Plus™ Poly(A) Polymerase Tailing Kit (CellScript, UK). The completed capping reaction was added directly to the poly(A)-tailing reaction (10x A-Plus poly(A) tailing buffer, 10 mM ATP and 20 units of A-Plus poly(A) polymerase). After incubation at 37 °C for 30 minutes, the reaction was stopped by ammonium acetate precipitation.

### Ammonium Acetate Precipitation

For the typical poly(A) tailing reaction, one volume of 5M ammonium acetate (ThermoFisher, UK) was added and mixed thoroughly with the reaction solution. After incubation on ice for 15 minutes, the RNA was pelletised by centrifuging it at 10,000x g for 15 minutes at 4°C. The RNA pellet was rinsed with 70% ethanol. After drying, the RNA pellet was resuspended in RNase-Free water.

### Agarose Gel Electrophoresis

Agarose was dissolved in 1x TAE buffer (0.75%, w/v) after being microwaved for 3 mins. The GelRed™ Nucleic Acid Gel Stain (Biotium, UK) was used to visualise the RNA.

For RNA gel electrophoresis, each sample was mixed with 2x RNA loading dye (New England Biolabs, UK). And the ssRNA ladder (New England Biolabs, UK) was used as a reference to estimate the size of the mRNA.

After loading ladders and samples to each well, the gel was run with a PowerPac™ universal power supply (Bio-Rad, UK) at 120 V for 1 hour. The image of the gel was acquired by UV lamp (UVITEC) with the associated software (UVI Platinum).

### Lipofectamine Transfection

Lipofectamine 2000 Reagent (Thermo Fisher Scientific, UK) was diluted in Gibco™ Opti-MEM™ medium (Thermo Fisher Scientific, UK). The mRNA samples (before and after poly(A) tailing and m^7^G capping) were then diluted in Gibco™ Opti-MEM™ and mixed with diluted Lipofectamine Reagent (1:1 ratio, final mRNA concentration: 0.25 μg/ml). The mRNA-Lipofectamine 2000 complexes were added to each well containing PEO1 cells. After 24 hours of incubation at 37 °C, the Bio-Glo™ Luciferase Assay System (Promega, UK) was added to each well and incubated in the dark for 15 minutes. Then the luminescence was measured by the Tecan Infinite M200 plate reader. Blank lipofectamine was used as control.

### Lipid Nanoparticle Synthesis

Different formulations of DOPE, DOTAP, DC-Chol·HCl and cholesterol (Avanti Polar Lipids, USA) **(Supplementary Table ST1)** were dissolved in chloroform with 1% (v/v) ethanol (lipid concentration: 10 mg/ml). The lipid solution was then concentrated by rotary evaporation at 40 °C under 350 mbar overnight, which offered a thin film on the wall of the flask. After evaporation, pre-warmed PBS (Sigma, UK) was used to hydrate the lipid film for 1-2 hours and produce an LNP suspension. The luciferase mRNA was then added to the suspension. To fix the size of the LNP, extrusion with a mini extruder (Avanti Polar Lipids, USA) was performed through a 0.1 or 0.2 μm polycarbonate membrane (Avanti Polar Lipids, USA) with up to 30 passes. The extrusion temperature was 50 °C, which exceeded the phase transition temperature (Tc) of all the lipids. LNPs synthesised by different lipid formulations have different Tc, below which it exists in a solid gel phase and above which a liquid crystalline phase.

### Cell Lines and Reagents

The PEO1, Kuramochi, OVCAR-4, and FT190 cell lines were supplied by American Type Culture Collection (ATCC). The cell lines were maintained in a 37 °C, 5% CO_2_ cell culture incubator using RPMI 1640 media (Sigma-Aldrich, UK) with 10% (v/v) Fetal Calf Serum (FCS, First Link, UK) and 1% (v/v) 10 mM L-Glutamine (Sigma-Aldrich, UK). Trypsin with 1 mM ethylenediaminetetraacetic acid (EDTA) was used to detach the cells from the flask.

### Characterisation of the LNPs

The size and zeta potential of the LNPs were tested by Malvern Zetasizer Nano ZS (Malvern Instruments, UK), and the data were analysed by Zetasizer Nano software (v3.30, Malvern Instruments Ltd).

The encapsulation efficiency of mRNA was analysed by Quant-it™ RiboGreen RNA Assay (Invitrogen, UK). Generally, LNP suspensions were diluted with 1x TE buffer (Invitrogen, UK) and 2% (v/v) Triton-100/TE buffer. Triton-100 was used to break the layers of phospholipids and release the encapsulated mRNA. After 10 minutes of incubation at 37 °C, the RiboGreen reagent was added to each sample, and the fluorescence was analysed by the Tecan Infinite M200 plate reader (Excitation: 485 nm; Emission: 528 nm, Gain: 55). The amount of mRNA in each sample was then quantified by comparing the fluorescence signal with the standard curve. The encapsulation efficiency (EE) was then calculated by the difference between the amount of mRNA in TE-diluted LNPs (A_1_) and TE/Triton-diluted LNPs (A_2_).

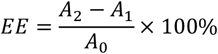

EE: Encapsulation efficiency.

A_1_: The amount of unencapsulated mRNA.

A_2_: The amount of total mRNA in the solution.

A_0_: The amount of mRNA added in the suspension before the extrusion.

### In vitro Transfection of Synthesised LNPs

The synthesised LNPs were used to transfect ovarian cancer cells in 96-well plates. Blank LNPs were used as controls. After the optimal treatment time (24 hours) **(Supplementary Figure SF3),** Bio-Glo™ Luciferase Assay System (Promega, UK) was added to each well and incubated in the dark for 15 minutes. The luminescence was measured by the Tecan Infinite M200 plate reader.

### Cytotoxicity analysis of LNPs

Cytotoxicity of LNPs was analysed through MTT (thiazolyl blue tetrazolium bromide) assay. MTT (Sigma-Aldrich, UK) was dissolved at a concentration of 10 % (v/v) in Dulbecco’s Modified Eagle Medium (high glucose, 4.5 g/L, Capricorn Scientific, UK). The MTT solution was then added to the cells after LNP treatment. After being incubated in a cell incubator (37 °C, 5% CO_2_) for 3 hours, the stop reagent (containing 10x sodium dodecyl sulphate (SDS) and 4 nM hydrochloric acid (HCl)) was added to each well to dissolve the blue formazan. The Tecan Infinite M200 plate reader was then used to measure the optical density at 570 nm of each well.

### Data Analysis

Statistical analysis was conducted using GraphPad Prism. Student’s two-sample t-tests, multiple t-tests and ANOVA tests were performed to analyse the statistical significance, which was indicated by asterisks: *p<0.05, **p<0.01, ***p<0.005, ****p<0.0001.

## Results

### mRNA Transcription and Modification

Luciferase mRNA was transcribed *in vitro*, as the input mRNA for downstream LNP optimisation experiments. The transcribed luciferase mRNA was around 1800 bases. After transcription, the obtained mRNA could be modified by adding a guanine nucleotide (“cap”) to the 5’ terminus (m^7^G capping), and adenosine monophosphates to the 3’-hydroxyl termini (poly(A) tailing) of the mRNA (*52*). After m^7^G capping and poly(A) tailing, the mRNA should become around 150 bp longer, so it is difficult to compare the length by agarose gel electrophoresis **(Figure 3A)**. To further verify whether the mRNA modification was successful, the transfection efficiency of lipofectamine loaded with these mRNAs were analysed. The lipofectamine loaded with luc-mRNA before poly(A) tailing and m^7^G capping showed no change compared to blank lipofectamine **(Figure 3B)**. However, the luminescence signal of lipofectamine loaded with luc-mRNA after poly(A) tailing and m^7^G capping was 50-fold higher than unmodified mRNA **(Figure 3B)**. As poly(A) tailing and m^7^G capping could improve the stability and translation efficiency of mRNA (*52*), the difference in luminescence signal indicated that luciferase mRNA was modified successfully.

**Figure 2.**
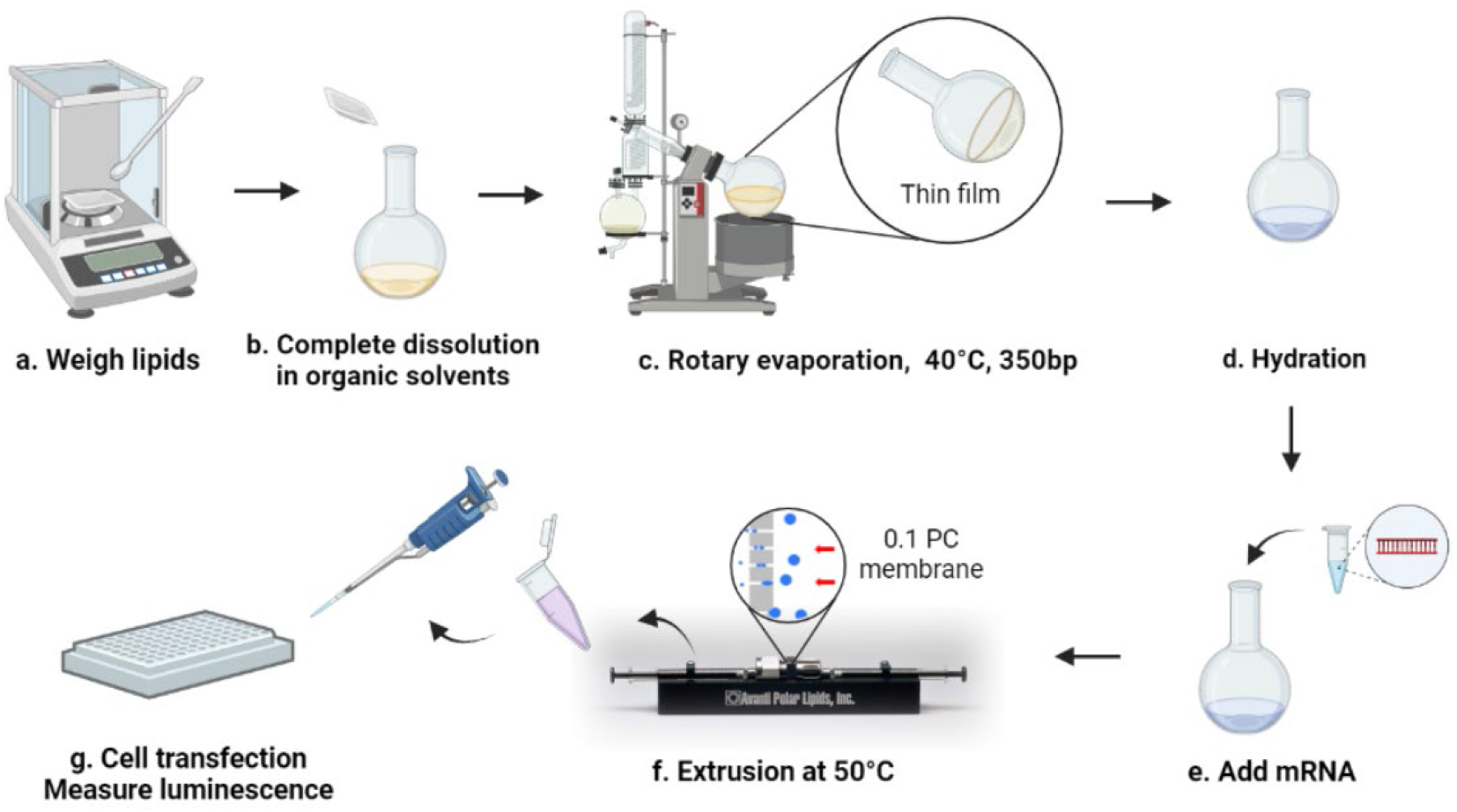
Method flow of the project.

**Figure 3.**
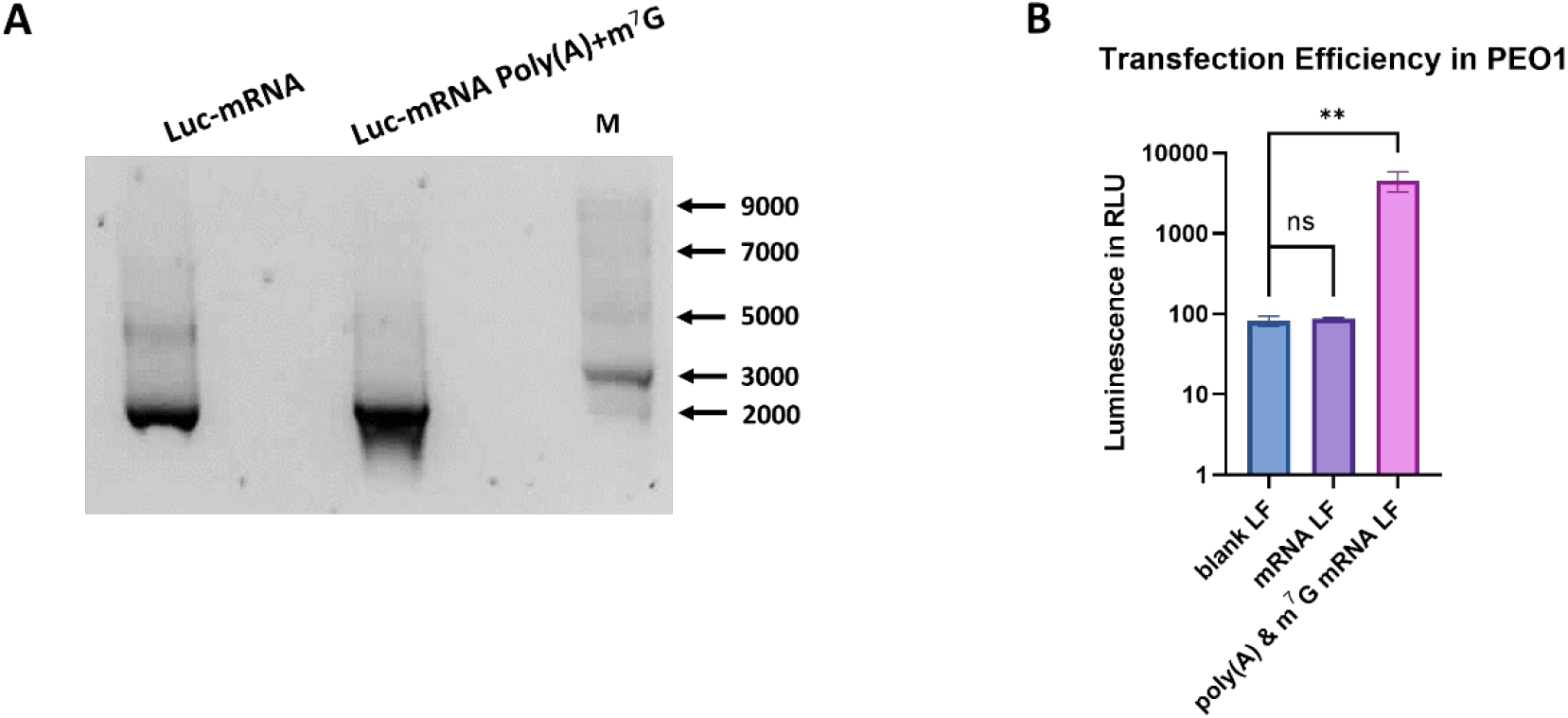
Message RNA transcription and modification. **(A)** RNA agarose gel electrophoresis of transcribed luciferase mRNA (Left Lane) and luciferase mRNA after poly(A) tailing and m^7^G capping (Middle lane). The amount of mRNA in each lane is 1 μg. Lane M contains the ssRNA ladder. The gel image shown is representative of 3 independent biological replicates. **(B)** Transfection efficiency of lipofectamine (LF) loaded with mRNA samples before and after poly(A) tailing and m^7^G capping. Luminescence signals are shown as the average relative light units (RLU) value ± standard deviation of 3 independent biological replicates. The two-tailed student t-tests are performed to analyse the statistical significance, which is indicated by asterisks: *p<0.05, **p<0.01.

### Optimisation of Lipid Formulations

Three different lipid formulations involving DOPE, DOTAP, DC-Chol, and cholesterol were used to synthesise LNPs **(Supplementary Table ST1)**. The molar ratio of cholesterol or DC-Chol in each formulation was 40% to ensure high LNP-cell fusion efficiency (*40*). The molar ratio of cationic/ionisable lipids was different in each formulation, which could affect the physical properties of the LNPs.

After the synthesis of LNPs using different formulations, their size distributions, zeta potential and encapsulation efficiency were analysed. The size distribution of DOPE/DC-Chol and DOPE/DOTAP/DC-Chol LNPs was narrow **(Figure 4A-B)**, of which the Polydispersity Index (PDI) was less than 0.1 **(Supplementary Table ST2)**, indicating good size homogeneity. However, the DOPE/DOTAP/Chol LNPs showed poor homogeneity (PDI > 0.2), with a wide size distribution from 100 to 1000 nm **(Figure 4C)**. For all three formulations, the average size of blank LNPs and mRNA-loaded LNPs were similar **(Figure 4D)**. The average size of DOPE/DC-Chol and DOPE/DOTAP/DC-Chol LNPs were around 129 nm, while the average size of DOPE/DOTAP/Chol LNPs was more than 500 nm **(Figure 4D)**.

**Figure 4.**
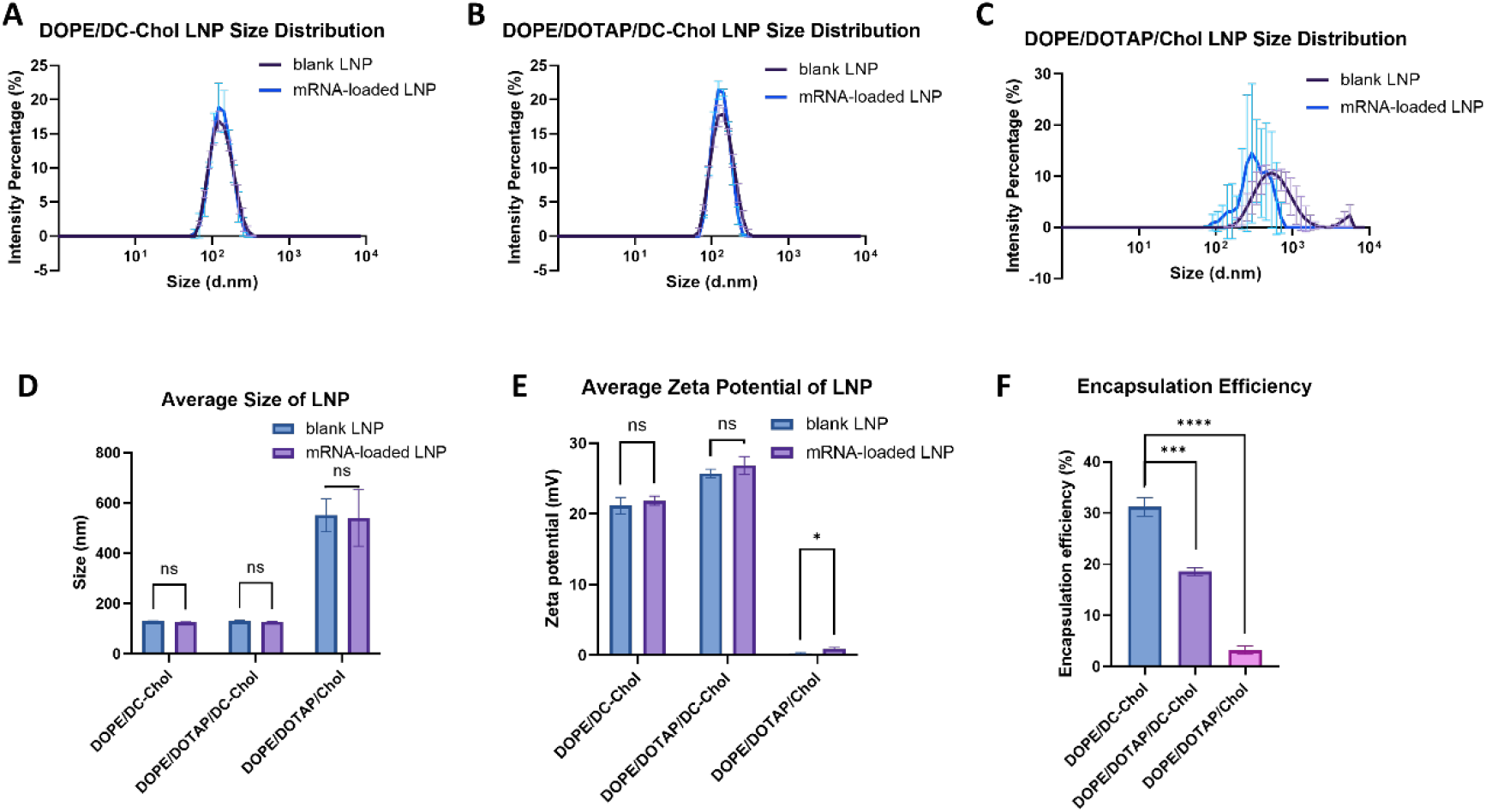
Characterisation of LNPs synthesised with different lipid formulations. **(A)** Size distribution of blank and luciferase mRNA-loaded DOPE/DC-Chol LNPs. **(B)** Size distribution of blank and luciferase mRNA-loaded DOPE/DOTAP/DC-Chol LNPs. **(C)** Size distribution of blank and luciferase mRNA-loaded DOPE/DOTAP/ Chol LNPs. **(D)** The average size of DOPE/DC-Chol, DOPE/DOTAP/DC-Chol, and DOPE/DOTAP/Chol LNPs. Results were analysed by multiple t-tests and ANOVA tests (p>0.05). **(E)** The average zeta potential of DOPE/DC-Chol, DOPE/DOTAP/DC-Chol, and DOPE/DOTAP/Chol LNPs. Results were analysed by multiple t-tests and ANOVA tests (p>0.05). **(F)** The encapsulation efficiency of DOPE/DC-Chol, DOPE/DOTAP/DC-Chol, and DOPE/DOTAP/Chol LNPs (analysed by students’ two-sample t-tests). Results are presented as the mean value ± standard deviation of 3 independent biological replicates. The statistical significance in t-tests is indicated by asterisks: *p<0.05, **p<0.01, ***p<0.005, ****p<0.0001. The input mRNA concentration before extrusion was 5 μg/ml. All the LNPs were extruded through a 0.1 μm Polycarbonate membrane for 30 passes.

For all three formulations, the difference between the average zeta potential of mRNA-loaded LNPs and blank LNPs was not significant **(Figure 4E)**. DOPE/DOTAP/DC-Chol LNPs were more cationic than DOPE/DC-Chol LNPs, with average zeta potential values of 26 mV and 21 mV, respectively **(Figure 4E)**. The average zeta potential of DOPE/DOTAP/Chol LNPs was around 0.5 mV, which was significantly lower than the other two formulations **(Figure 4E)**.

DOPE/DC-Chol LNPs had the best mRNA encapsulation efficiency, which was 31.2% **(Figure 4F)**. The encapsulation efficiency of DOPE/DOTAP/DC-Chol was 18.6%, followed by DOPE/DOTAP/Chol LNPs, with an encapsulation efficiency of 3.2% **(Figure 4F)**.

Overall, DOPE/DC-Chol and DOPE/DOTAP/DC-Chol LNPs demonstrated narrow size distributions, appropriate size ranges and relatively high mRNA encapsulation efficiencies. In comparison, due to wide size distribution and low encapsulation efficiency, DOPE/DOTAP/Chol LNPs will not be used for further optimisation.

### Optimisation of input mRNA Concentration

As DOPE/DC-Chol and DOPE/DOTAP/DC-Chol LNPs were selected for further optimisation, the influence of input mRNA concentration (IRC) before extrusion on encapsulation and transfection efficiency of these two kinds of LNPs was investigated. The size distribution of DOPE/DC-Chol and DOPE/DOTAP/DC-Chol LNPs with different IRCs were narrow **(Figure 5A-B)**, where the Polydispersity Indexes (PDIs) were all below 0.1 **(Supplementary Table ST2)**.

**Figure 5.**
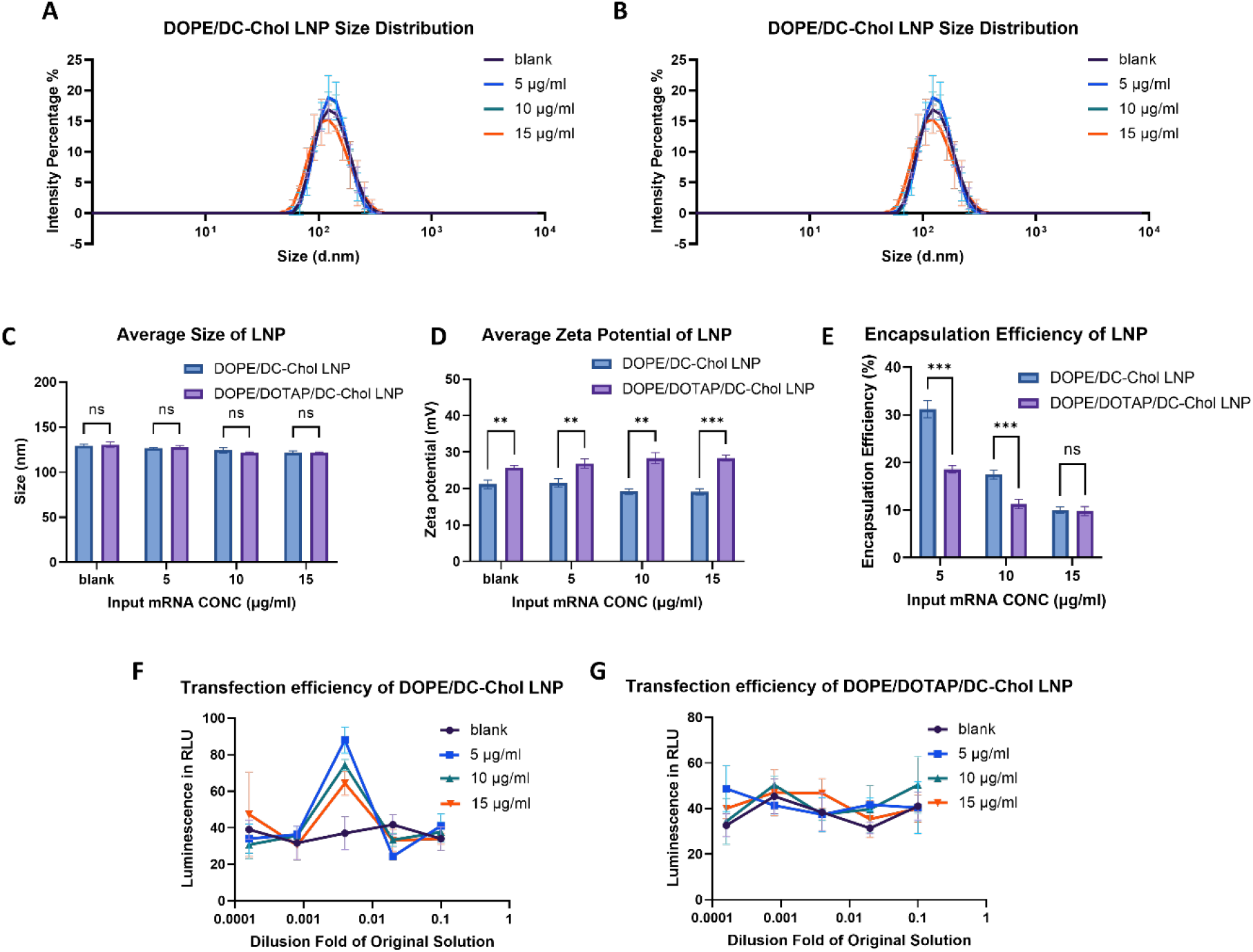
Characterisation of DOPE/DC-Chol and DOPE/DOTAP/DC-Chol LNPs with different input mRNA concentrations (IRCs). **(A)** Size distribution of DOPE/DC-Chol LNPs with IRCs of 5, 10 and 15 μg/ml. **(B)** Size distribution of DOPE/DOPTAP/DC-Chol LNPs with IRCs of 5, 10 and 15 μg/ml. **(C)** The average size of DOPE/DC-Chol and DOPE/DOPTAP/DC-Chol LNPs with IRCs of 5, 10 and 15 μg/ml. Results are analysed by multiple t-tests and ANOVA test (p>0.05). **(D)** The average zeta potential of DOPE/DC-Chol and DOPE/DOPTAP/DC-Chol LNPs with IRCs of 5, 10 and 15 μg/ml. Results are analysed by multiple t-tests and ANOVA test (p<0.01). **(E)** The encapsulation efficiency of DOPE/DC-Chol and DOPE/DOPTAP/DC-Chol LNPs with IRCs of 5, 10 and 15 μg/ml. Results are analysed by multiple t-tests and ANOVA test (p<0.0001). **(F)** The transfection efficiency of DOPE/DC-Chol LNPs with different IRCs in the PEO1 cell line. Blank DOPE/DC-Chol LNPs are used as control. **(G)** The transfection efficiency of DOPE/DOTAP/DC-Chol LNPs with different IRCs in the PEO1 cell line. Blank DOPE/DOTAP/DC-Chol LNPs are used as control. Results are demonstrated as the mean value ± standard deviation of 3 independent biological replicates. The statistical significance in multiple t-tests is indicated by asterisks: *p<0.05, **p<0.01, ***p<0.005, ****p<0.0001. All the LNPs were extruded through a 0.1 μm Polycarbonate membrane for 30 passes.

For both formulations, the change of input mRNA concentration did not have a significant effect on the average size and zeta potential of the LNPs **(Figure 5C-D)**. However, the zeta potential of DOPE/DC-Chol and DOPE/DOTAP/DC-Chol LNPs demonstrated a more significant difference as the IRC increased **(Figure 5D)**. The encapsulation efficiency for both DOPE/DC-Chol and DOPE/DOTAP/DC-Chol LNPs decreased as the IRC increased **(Figure 5E)**. At all IRCs, DOPE/DC-Chol LNPs had higher encapsulation efficiency than DOPE/DOTAP/DC-Chol LNPs **(Figure 5E)**.

The transfection efficiency of the two types of LNPs with different IRCs was analysed in the PEO1 cell line. For DOPE/DC-Chol LNPs, the highest luminescence signal could only be achieved when the dilution fold (DF) was 0.004. The luminescence signal was close to control when the DF value was above or below 0.004. **(Figure 5F)**. Among all the IRCs analysed, a lower IRC could result in better transfection efficiency of DOPE/DC-Chol LNPs **(Figure 5F)**. However, for DOPE/DOTAP/DC-Chol LNPs, the luminescence signal of mRNA-loaded LNPs did not change compared to blank LNPs, indicating very low transfection efficiency **(Figure 5G)**.

LNPs with IRC less than 5 μg/ml did not achieve higher encapsulation and transfection efficiency **(Supplementary Figure SF4)**. Therefore, DOPE/DC-Chol LNPs with an IRC of 5 μg/ml, which showed the best encapsulation efficiency **(31.2%)** and transfection efficiency **(1.4-fold of blank LPS)** among all the formulations with different IRCs, were selected for further optimisation.

### Optimisation of Extrusion Process

The effect of the LNP extrusion process on the encapsulation and transfection efficiency was then investigated. The LNPs synthesised all had high size homogeneity, with PDI values of less than 0.1 **(Supplementary Table ST2)**. The size distribution of DOPE/DC-Chol LNPs extruded through a 0.1 μm PC membrane was narrower than that extruded with a 0.2 μm PC membrane **(Supplementary Table ST2)**. Increasing extrusion passes resulted in a more narrow size distribution **(Figure 6A)**. LNPs extruded with 0.2 μm PC membrane had an average diameter of 141 nm, which was about 10-20 nm more than the average size of LNPs extruded with 0.1 μm PC membrane **(Figure 6B)**.

**Figure 6.**
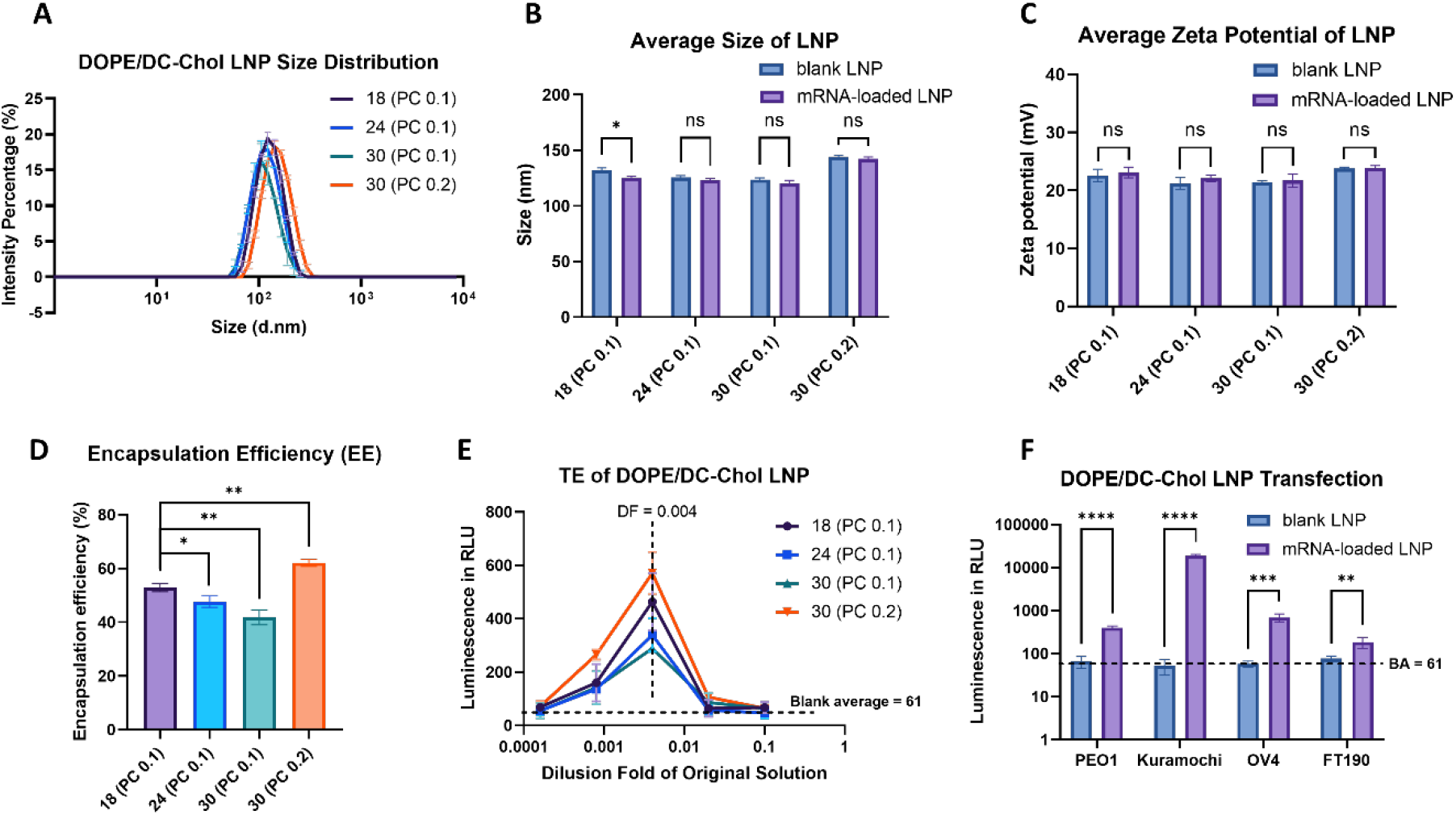
Characterisation of DOPE/DC-Chol LNPs with different extrusion methods. **(A)** The size distribution of DOPE/DC-Chol LNPs extruded through a 0.1 μm PC membrane for 18, 24, and 30 passes and LNPs extruded through a 0.2 μm PC membrane for 30 passes. **(B)** The average size of DOPE/DC-Chol LNPs with different extrusion methods. Results are analysed by multiple t-tests and ANOVA test (p>0.05). **(C)** The average zeta potential of DOPE/DC-Chol LNPs with different extrusion methods. Results are analysed by multiple t-tests and ANOVA test (p>0.05). **(D)** The encapsulation efficiency of DOPE/DC-Chol LNPs with different extrusion methods (analysed by students’ two-sample t-tests). **(E)** The transfection efficiency of DOPE/DC-Chol LNPs with different extrusion methods in PEO1 cell line. The transfection efficiency of the blank LNPs is represented by a baseline indicating their average signal. **(F)** The transfection luminescence signal of DOPE/DC-Chol LNPs in PEO1, Kuramochi, OVCAR-4 and FT190 cell lines. Results are analysed by multiple t-tests and ANOVA test (p<0.0001). The LNPs are extruded through a 0.2 μm PC membrane for 30 passes, and the dilution fold of the used LNPs is 0.004. Results are demonstrated as the mean value ± standard deviation of 3 independent biological replicates. The statistical significance in t-tests is indicated by asterisks: *p<0.05, **p<0.01, ***p<0.005, ****p<0.0001. The input mRNA concentration before extrusion was 5 μg/ml.

For LNPs extruded with 0.1 μm PC membrane, the zeta potential showed minimal changes under different extrusion passes **(Figure 6C)**. The difference between the zeta potential of LNPs extruded with 0.2 μm PC membrane and that extruded with 0.1 μm PC membrane was non-significant **(Figure 6C)**.

The encapsulation efficiency of LNPs also showed similar trends as the zeta potential. For LNPs extruded with 0.1 μm PC membrane, the encapsulation efficiency also decreased as the extrusion passes increased **(Figure 6D)**. LNPs extruded with 0.2 μm PC membrane had higher encapsulation efficiency than that extruded with 0.1 μm PC membrane **(Figure 6D)**. The transfection efficiency of the LNPs was highly correlated with their encapsulation efficiency **(Figure 6E)**: The DOPE/DC-Chol LNPs extruded with 0.2 μm PC membrane for 30 passes showed the best encapsulation efficiency **(62%)** and transfection efficiency **(9.4-fold RLU of baseline) (Figure 6D-E),** thus deemed as the optimised LNP under the conditions used.

This optimised LNP was then tested in a panel of ovarian cancer cell lines (PEO1, Kuramochi, OVCAR-4) and one non-malignant cell line (FT190). Kuramochi showed the highest luminescence signal (307.4-fold RLU of baseline), followed by OVCAR-4 (11.3-fold RLU of baseline) and PEO1 (9.4-fold RLU of baseline) **(Figure 6F)**. FT190 showed the lowest transfection luminescence signal, which was only 3-fold RLU of baseline (**Figure 6F)**.

### Stability and Biocompatibility of optimised DOPE/DC-Chol LNPs

The sizes of blank and mRNA-loaded LNPs were relatively stable over 14 days after synthesis, though there was a decrease by 1 nm for both LNPs **(Figure 7A)**. The zeta potential of blank DOPE/DC-Chol stayed at around 21.5 mV during the 14 days **(Figure 7B)**. In comparison, the zeta potential of mRNA-loaded LNPs started to decrease from day 4 and kept declining afterwards **(Figure 7B)**, which suggests the leakage of the negatively charged mRNA molecule. The zeta potential of mRNA-loaded LNPs was higher than blank LNPs before day 6, after which the mRNA-loaded LNPs became less cationic than the blank LNPs **(Figure 7B)**. The mRNA encapsulation efficiency decreased by 30% on day 7 and dropped to around 5% on day 14 **(Figure 7C)**. Regarding the transfection efficiency, the luminescence signal of the day-7 LNPs’ transfection was less than half of that for the day-1 LNP **(Figure 7D)**. The luminescence signal of the day-14 LNP transfection dropped to less than one-sixth of that for freshly prepared LNPs, which was around 1.6-fold of blank LNP **(Figure 7D)**.

**Figure 7.**
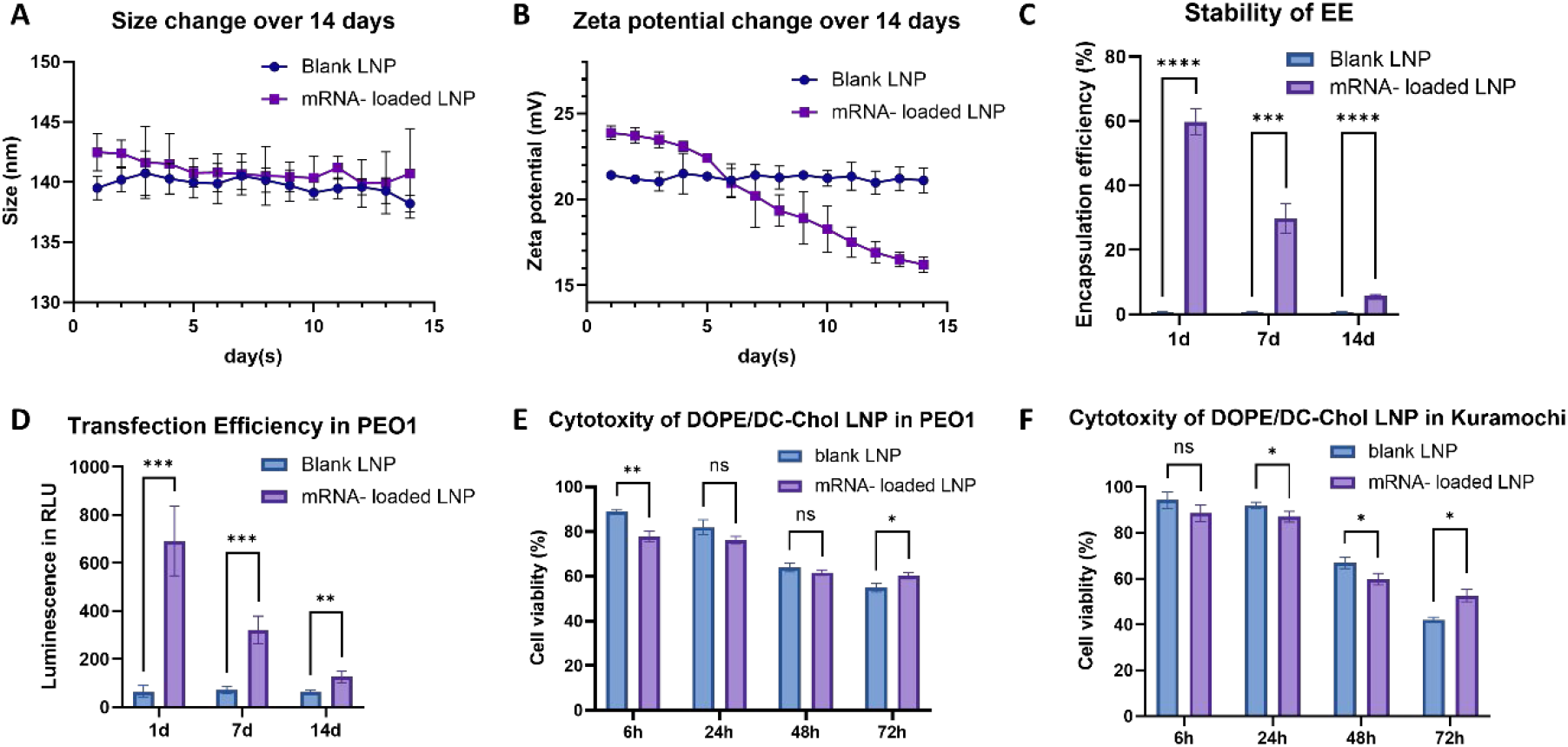
Stability and biocompatibility of the optimised DOPE/DC-Chol LNPs. **(A)** Size change of blank LNP and mRNA-loaded LNP over 14 days. **(B)** Zeta potential change of blank LNP and mRNA-loaded LNP over 14 days. **(C)** Encapsulation efficiency of blank LNP and mRNA-loaded LNP kept 1 day, 7 days and 14 days after synthesis. Results are analysed by multiple t-tests and ANOVA test (p<0.0001). **(D)** Transfection efficiency in PEO1 cell line of blank LNP and mRNA-loaded LNP kept for 1 day, 7 days and 14 days after synthesis. Results are analysed by multiple t-tests and ANOVA test (p<0.005). **(E)** Cell viability of PEO1 and Kuramochi cells after being treated by freshly made blank LNPs for 6, 24, 48 and 72 hours. Results are analysed by multiple t-tests and ANOVA test (p>0.05). **(F)** Cell viability of PEO1 and Kuramochi after being treated by freshly made mRNA-loaded LNPs for 6, 24, 48 and 72 hours. Results are analysed by multiple t-tests and ANOVA test (p>0.05). Input RNA concentration: 5 μg/ml; Pore diameter of PC membrane: 0.2 μm; Dilution fold: 0.004. The results are demonstrated as the average value ± standard deviation of three independent biological replicates. The statistical significance in multiple t-tests is indicated by asterisks: *p<0.05, **p<0.01, ***p<0.005, ****p<0.0001.

The cell viability decreased when the LNP treatment time was increased in both PEO1 **(Figure 7E)** and Kuramochi cells **(Figure 7F)**. The cell viability was above 75% within 24 hours of treatment, after which it declined rapidly **(Figure 7E-F)**. The difference between the sensitivity of PEO1 and Kuramochi cells towards the LNPs was not significant **(Figure 7E-F)**. Also, there was little biological difference between the two LNPs. However, statistically, mRNA-loaded LNPs were slightly more cytotoxic than blank LNPs within 48 hours of treatment **(Figure 7E-F)**.

## Discussion

In this project, we synthesised different types of cationic LNPs, and optimised their encapsulation and transfection efficiency by adjusting the lipid formulation, input mRNA concentration and extrusion process. After optimisation, the encapsulation efficiency was successfully improved to 62%, resulting in a high transfection luminescence signal (9.4 times compared to baseline).

We first transcribed luciferase mRNA and modified it by adding poly(A) tail and m^7^G cap. Then we used this luciferase mRNA as a reporter to evaluate the encapsulation and transfection efficiency as we optimised the synthesised LNPs.

In the lipid formulation optimisation, we analysed the size, zeta potential and encapsulation efficiency of DOPE/DC-Chol, DOPE/DOTAP/DC-Chol, and DOPE/DOTAP/Chol-formulated LNPs. We found that DOPE/DC-Chol and DOPE/DOTAP/DC-Chol LNPs had better size homogeneity and mRNA encapsulation efficiency. DOPE/DOTAP/Chol LNPs had the poorest size distribution and encapsulation efficiency. The average size of DOPE/DOTAP/Chol LNPs was around 550 nm. However, a 0.1 PC membrane will usually produce LNPs with a much smaller size (100-150 nm) (*53*). This could be because this formulation included the lowest dose of cationic/ionisable lipids, and the zeta potential was very low, which might result in LNP agglomeration (*54*). The agglomerated LNPs could expand or clog (*55*) the pore diameter of the PC membrane during extrusion. In addition, due to the low ratio of cationic/ionisable lipids, the electrostatic interaction between lipids and negatively charged mRNA molecules was very weak, resulting in lower mRNA encapsulation efficiency (*56*). However, although DOPE/DOTAP/DC-Chol LNPs were more cationic than DOPE/DC-Chol LNPs, the encapsulation efficiency was still lower. This could be due to the increased concentration of DOPE, which has a pronounced cone shape compatible with hexagonal HII organisation, providing improved packing properties (*57*).

During the optimisation of input mRNA concentration, we found that DOPE/DC-Chol LNPs with an IRC of 5 μg/ml showed the best encapsulation and transfection efficiency. The IRC did not have a significant effect on the size distribution and zeta potential of the LNPs. However, the encapsulation efficiency was found to be negatively correlated with the IRC. This might suggest that there could be a limit to the maximum amount of mRNA that can be encapsulated, depending on the flexibility and properties of different LNPs (*58*). The transfection efficiency of the LNPs was positively correlated with the encapsulation efficiency instead of the amount of mRNA encapsulated, which was also verified by other studies (*59*, *60*). The transfection efficiency of DOPE/DOTAP/DC-Chol LNPs was very low, although it showed similar encapsulation efficiency as DOPE/DC-Chol LNPs at certain IRCs. This may be explained by the excessively strong electrostatic interaction between lipids and mRNA molecules. Studies have shown that when the LNPs are too cationic, it can be difficult for mRNA molecules to detach from the lipids. This could be improved by reducing the ratio of cationic/ionisable lipids in the formulation (*61*).

During the extrusion optimisation, DOPE/DC-Chol LNPs extruded with a 0.2 μm PC membrane for 30 passes showed the best encapsulation and transfection efficiency. These results suggested that more extrusion passes and a smaller diameter of the PC membrane could lead to better homogeneity, but lower encapsulation efficiency of the LNPs. The effect on encapsulation efficiency could be explained by the rationale of the extrusion process. During extrusion, the lipid layers will break and form smaller vesicles, during which the LNPs could encapsulate mRNA molecules. However, there is a threshold where the amount of encapsulated mRNA becomes less than the amount of released mRNA caused by the break-and-form process (*62*). Therefore, it is important to ensure that the extrusion process produces LNPs with a reasonable level of size homogeneity and encapsulation efficiency simultaneously. After a series of optimisation approaches, we finally improved the encapsulation efficiency to 62% and the transfection RLU signal in PEO1 cells to 9.4-fold of baseline.

Further, we found that the transfection luminescence signal of the optimised LNP was higher in ovarian cancer cell lines than in the non-malignant cell line FT190, which might be caused by the biophysical properties of different cell lines (i.e. membrane flexibility, internalisation ability, etc.) (*63*).

Finally, we analysed the stability and biocompatibility of the optimised LNPs. The amount of leaked mRNA increased as the preservation time increased, resulting in decreased zeta potential, encapsulation efficiency and transfection efficiency over 14 days. The cytotoxicity of the LNPs in PEO1 and Kuramochi cells increased as the duration of the treatment lengthened. This could be because the membrane function and integrity of the cell or subcellular compartments were affected when exposed to cationic lipids for longer periods (*63*). Despite this, the LNPs showed relatively high stability and biocompatibility within 24 hours. This suggests that the LNPs should be freshly synthesised for each use.

Admittedly, this project has a range of limitations due to time constraints. First, as for the modification of luciferase mRNA, we used an indirect way to analyse whether the poly(A) tailing and m^7^G capping were successful. More accurate ways like CapQuant (*64*) and RNA sequencing platforms (e.g. Pacific Biosciences and Oxford Nanopore Technologies) (*65*) could be applied to perform the quality control of the modification. Second, regarding the lipid formulation optimisation, we only included 4 different kinds of lipids. More lipid formulations could be tested, for example, other widely used lipids like DODAC (*66*) and DSPC (*67*) could be included at different dosage ratios. In this case, further adjustment of the zeta potential of the LNPs might be done to achieve higher encapsulation and transfection efficiency. Third, regarding the optimisation of the extrusion process, high-pressure extrusion with an automatic extruder can ensure better homogeneity and higher yields than the manual process (*68*, *69*). In addition, as for the characterisation of the LNPs, cryo-electron microscopy (cryoEM) could be used to visualise the structure of the LNPs and identify whether the LNPs are multi-layered, in order to evaluate the efficiency of the extrusion process (*70*, *71*). Besides, the transfection of the optimised LNPs was carried out only in a limited range of cell lines. More ovarian cancer cell lines like OVCAR-3 and PEA1, and more non-malignant cell lines could be included to compare the transfection luminescence signal. If more evidence suggests the luminescence signal of the optimised LNPs is higher in ovarian cancer cell lines than in non-malignant cell lines, it might be possible to utilise the difference to achieve targeted mRNA delivery to ovarian cancer cell lines. Studies have shown that LNPs could be customised as targeted delivery systems without additional targeting compounds attached (*72*, *73*). For example, Kranz et al. used DOTMA/DOPE LNPs with a specific lipid component ratios and IRC to target dendritic cells for cancer immunotherapy (*72*). Finally, when analysing the stability of the LNPs, we could also test the phase transition temperature of the LNPs by differential scanning calorimetry (DSC) to measure their thermal stability (*74*, *75*).

For future work, surface modification of the optimised LNPs could further increase their stability and targeting efficiency. The lipids could be modified with PEG_2000_ before the LNP synthesis, as PEGlyted cationic LNPs have been proved to have higher stability, better biocompatibility and biodistribution when used *in vivo* (*76*, *77*). Apart from PEG, specific groups like alkynes could be linked to the lipids as a “click” handle for reaction with an azide. This is a simple and wildly applied ligand fixation method with extremely high yield and tolerance of functional groups (*78*). Our group has successfully linked an azide to folic acid (FA). Therefore, through the “click” reaction, the FA-modified LNP could be synthesised to target folate receptors (FOLR), which are overexpressed in most ovarian cancer cell lines (*79*). Furthermore, we could attach ^18^F onto the LNP surface for positron emission tomography (PET) imaging (*80*), making the LNP a multifunctional platform for cancer theranostics.

Our lab is currently setting up the protocol of synthesising self-amplifying IL-15 mRNA, which is closely related to the immunotherapy response of ovarian cancer (*81*, *82*). After the establishment of the complete mRNA delivery system, the luciferase reporter could be replaced by the self-amplifying IL-15 mRNA. The theranostic platform will then be analysed in 3D tumour models or even *in vivo* to determine whether it could improve the local immune response and prognosis in ovarian cancer.

In conclusion, this project optimised the mRNA encapsulation and transfection efficiency of DOPE/DC-Chol cationic LNPs by adjusting the different reaction parameters. Apart from the relatively high encapsulation and transfection efficiency, the LNPs we developed also had stable physical characteristics and high biocompatibility within 24 hours. These optimised LNPs could be further modified with targeting and/or imaging agents, which could serve as a potential theranostic platform for ovarian cancer immunotherapy.

## Abbreviations

ANOVA: analysis of variance
ATCC: American Type Culture Collection
ATP: adenosine triphosphate
CryoEM: cryoelectron microscopy
CTP: cytidine triphosphate
DC-Chol: 3β-[N-(N’,N’-dimethylaminoethane)-carbamoyl] cholesterol
DF: dilution fold
DNA: Deoxyribonucleic acid
DNase: deoxyribonuclease
DOPE: 1,2-dioleoyl-sn-glycero-3-phosphoethanolamine
DOTAP: 1,2-dioleoyl-3-trimethylammonium-propane
DOTMA: 2-di-O-octadecenyl-3-trimethylammonium-propane
DSC: differential scanning calorimetry
EDTA: ethylenediaminetetraacetic acid
EE: encapsulation efficiency
FA: folic acid
FCS: fetal calf serum
FOLR: folate receptor
GTP: guanosine triphosphate
IRC: input RNA concentration
IVT: *in vitro* transcription
LF: lipofectamine
LNP: lipid nanoparticle
Luc: luciferase
mRNA: messager ribonucleic acid
MTT: thiazolyl blue tetrazolium bromide
OVA: ovalbumin
PC: polycarbonate
PDI: polydispersity index
PEG: polyethylene glycol
PET: positron emission tomography
RLU: relative light units
RNase: Ribonucleases
SAM: S-adenosyl-methionine
SDS: sodium dodecyl sulfate
ssRNA: single-stranded RNA
TAE: Tris-acetate-EDTA
TE: transfection efficiency
UTP: uridine triphosphate
UV: ultraviolet

