## Supplementary Materials for "Development and optimisation of cationic lipid nanoparticles for mRNA delivery"

### Supplementary Methods *(Preliminary experiments of IL-15 mRNA preparation)*

#### *IL-15 DNA Restriction Enzyme Digestion and Clean-up*

IL-15 DNA template was supplied by GenScript Biotech (UK). One unit of NruI restriction enzyme was used to digest 1 µg of DNA template in 1 hour at 37 °C in a total reaction volume of 50 µl. The DNA was diluted by 1x NEBuffer™ r3.1 (New England Biolabs, UK). The digested DNA was mixed with Wizard® DNA clean-up resin (Promega, UK) and pushed through Wizard® DNA clean-up minicolumn (Promega, UK). After being washed with isopropanol, the linear DNA was detached from the column by pre-warmed (65-70 °C) RNase-free water.

#### *IL-15 DNA Agarose Gel Electrophoresis*

Agarose was completely dissolved in 1x TAE buffer (0.75%, w/v) after being microwaved for 3 mins. The 10,000x GelRed™ Nucleic Acid Gel Stain (Biotium, UK) was added when the agarose solution cooled down to about 50 °C. The solution was poured into a gel tray with a comb and placed at room temperature to solidify. The gel was then placed in a gel box filled with 1x TAE buffer. For DNA gel electrophoresis, each sample containing 1 µg of cleaned linear DNA was mixed with 5x DNA loading dye (Bioline, UK). And the GeneRuler 1kb Plus DNA ladder (ThermoFisher, UK) was used as a reference to estimate the size of sample DNA.

After loading ladders and samples to each well, the gel was run with a PowerPac™ universal power supply (Bio-Rad, UK) at 120 V for 1 hour. The image of the gel was acquired by UV lamp (UVITEC) with the associated software (UVI Platinum).

### Supplementary Figures

#### *Lipid Formulation in LNP Synthesis*

| | Formulation | DOPE (mM) | DC-Chol (mM) | DOTAP (mM) | Chol (mM) |
|---|---|---|---|---|---|
| 1 | DOPE/DC-Chol | 15 | 10 | 0 | 0 |
| 2 | DOPE/DOTAP/DC-Chol | 7.5 | 10 | 7.5 | 0 |
| 3 | DOPE/DOTAP/Chol | 7.5 | 0 | 7.5 | 10 |

***Supplementary Table ST1.*** **Lipid formulations involved in the optimisation.** The dosage ratio of cholesterol and cholesterol-derived lipids is 40% in each formulation.

#### *Polydispersity Index (PDI) of the LNPs*

| Formulation | IRC (µg/ml) | Extrusion Passes | PC membrane (µm) | PDI |
|---|---|---|---|---|
| DOPE/DC-Chol | 0 | 30 | 0.1 | 0.085 |
| | 2.5 | 30 | | 0.099 |
| | 5 | 10 | 0.1 | **0.211** |
| | | 13 | | **0.182** |
| | | 15 | | **0.137** |
| | | 18 | | 0.083 |
| | | 24 | | 0.077 |
| | | 30 | | 0.068 |
| | 5 | 10 | 0.2 | **0.193** |
| | | 13 | | **0.174** |
| | | 15 | | **0.171** |
| | | 18 | | **0.148** |
| | | 24 | | **0.129** |
| | | 30 | | 0.079 |
| | 10 | 30 | 0.1 | 0.047 |
| | 15 | 30 | | 0.036 |
| DOPE/DOTAP/DC-Chol | 0 | 30 | 0.1 | 0.091 |
| | 5 | | | 0.069 |
| | 10 | | | 0.056 |
| | 15 | | | 0.063 |
| DOPE/DOTAP/Chol | 0 | 30 | 0.1 | **0.351** |
| | 5 | | | **0.385** |

***Supplementary Table ST2.*** **Polydispersity Index (PDI) of the synthesised LNPs.** The LNPs with PDI > 0.1 were not included in the optimisation process, which is highlighted in bold font. The PDI values shown are the average value of three technical replicates.

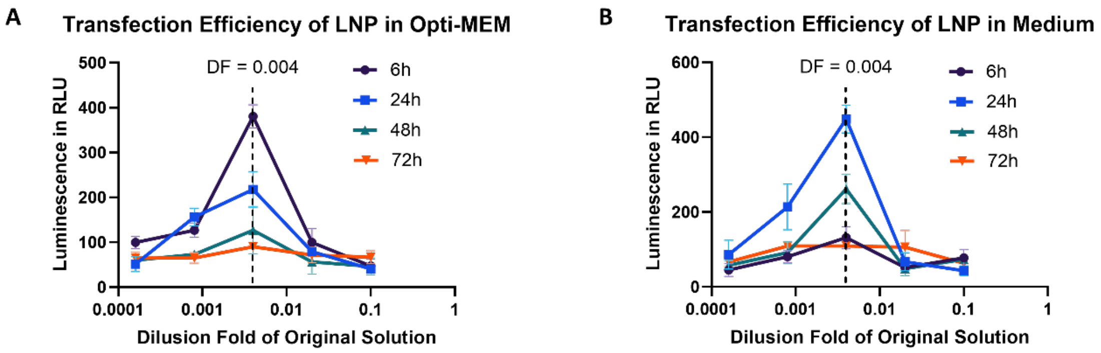


***Supplementary Figure SF3.*** **Transfection efficiency of DOPE/DC-Chol LNP in medium and opti-MEM for different treatment duration. (A)** The transfection signal of DOPE/DC-Chol LNPs in the opti-MEM after 6, 24, 48 and 72 hours of treatment. **(B)** The transfection signal of DOPE/DC-Chol LNPs in the medium after 6, 24, 48 and 72 hours of treatment. Results are demonstrated as the mean value ± standard deviation of 3 independent biological replicates.

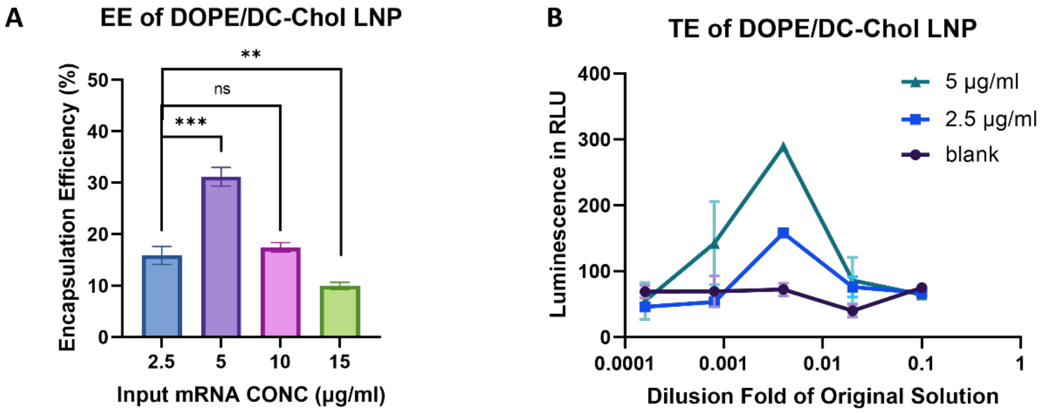


***Supplementary Figure SF4.*** **Encapsulation and transfection efficiency of LNPs with different IRCs. (A)** The encapsulation efficiency of DOPE/DC-Chol LNPs with IRCs of 2.5, 5, 10, and 15 μg/ml. **(B)** The transfection efficiency of DOPE/DC-Chol LNPs with IRCs of 2.5 and 5 μg/ml in PEO1 cell lines. Results are demonstrated as the mean value ± standard deviation of 3 independent biological replicates. Multiple t-tests are performed to analyse the statistical significance, which is indicated by asterisks: *p<0.05, **p<0.01, ***p<0.005. LNPs were extruded through a 0.1 μm Polycarbonate membrane for 30 passes.

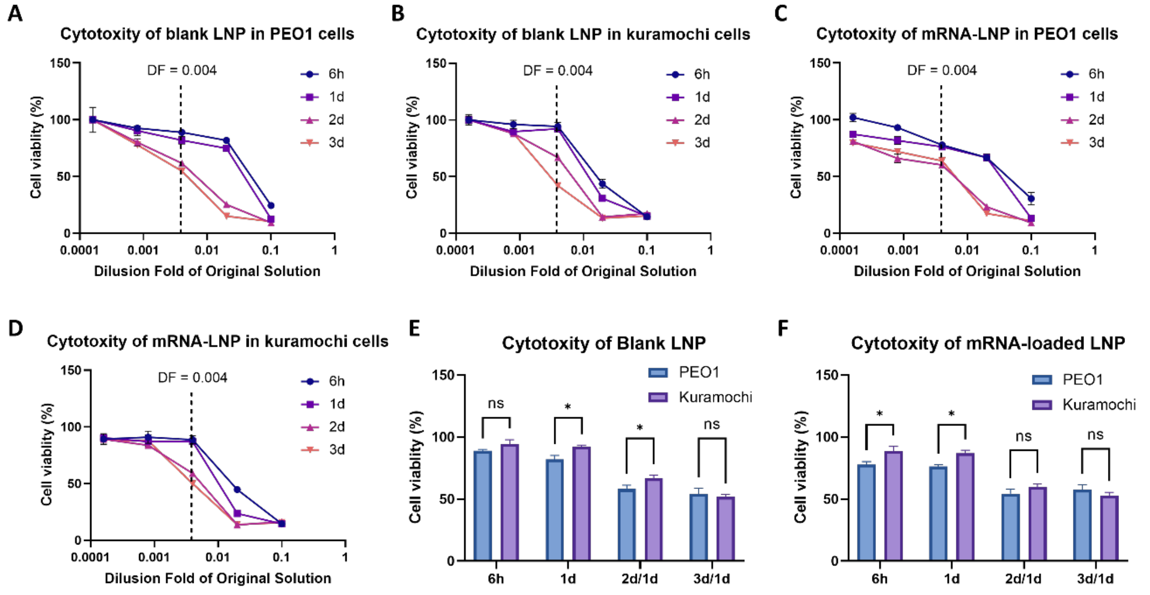


***Supplementary Figure SF5.*** **Cytotoxicity of the optimised DOPE/DC-Chol LNPs in PEO1 and Kuramochi cells. (A)** Cell viability of PEO1 cells after being treated by blank LNPs with different DFs for 6, 24, 48 and 72 hours. **(B)** Cell viability of Kuramochi cells after being treated by blank LNPs with different DFs for 6, 24, 48 and 72 hours. **(C)** Cell viability of PEO1 cells after being treated by mRNA-loaded LNPs with different DFs for 6, 24, 48 and 72 hours. **(D)** Cell viability of Kuramochi cells after being treated by mRNA-loaded LNPs with different DFs for 6, 24, 48 and 72 hours. **(E)** Cell viability of PEO1 and Kuramochi after being treated by blank

LNPs. The medium was changed every 24 hours after being treated for one day. **(F)** Cell viability of PEO1 and Kuramochi after being treated by mRNA-loaded LNPs. The medium was changed every 24 hours after being treated for one day.
